# Emergence of multiple collective motility modes in a physical model of cell chains

**DOI:** 10.1101/2025.01.30.635787

**Authors:** Ying Zhang, Effie E. Bastounis, Calina Copos

## Abstract

Collective cell migration underpins key (patho)physiological processes, ranging from embryonic development to wound healing and cancer metastasis. While notable progress has been made in elucidating mechanisms that drive collective cell motility, the classification remains incomplete. In this study, we focus on the migration patterns of small cell chains, specifically cohesive pairs of cells migrating after each other on flat surfaces. Experiments with *Dictyostelium discoideum* (Dd) cells, which typically display amoeboid motility, revealed two distinct motility modes in cell pairs: the individual contributor (IC) mode, where each cell generates its own traction force dipole, and the supracellular (S) mode, characterized by a single dipole. Intriguingly, the IC mode dominates in Dd pairs, but the S mode prevails in Madin-Darby canine kidney (MDCK) cell doublets, which typically undergo mesenchymal motility. This observation highlights an apparent discrepancy in emergent motility modes between cell types. To uncover the physical mechanisms driving these diverse motility modes, we developed a two-dimensional biophysical model incorporating mechanochemical details such as cell-cell adhesion, combined with membrane-cortex contractility, and cell-matrix adhesion. Our model could recapitulate many experimental observations; the IC mode emerged naturally in amoeboid doublets when both cells exerted similar traction stresses, while the S mode dominated with “stronger” leaders that essentially pull on trailers. In contrast, in our simulations, mesenchymal MDCK-like pairs largely migrated in supracellular arrangement (S mode), with traction stress patterns representative of a rear-drive system with a “pushy” trailer, rather than a front-drive system. Our findings also showed that increasing cell-matrix adhesion predisposes amoeboid cell chains to act autonomously (IC mode), but the chain’s motility mode was largely insensitive to changes in cell-cell adhesion parameters. Contrary to amoebas, MDCK-like cell chains showed a bias towards S mode when increasing cell-matrix adhesion and a preference on IC mode when increasing cell-cell adhesion. Extending the model to longer cell chains, we showcase the model’s applicability across scales, providing a foundation for exploring collective migratory behavior in other contexts.

## I. INTRODUCTION

Living cells are able to mechano-sense their local environment and respond by adapting their cytoskeleton, biomechanics, and motility [1–3]. At the single cell level, the motility mechanism can range from mesenchymal [4, 5], gliding [6], stepping-like [7, 8], to bleb-based migration in low adhesive environments [9, 10] and even rotational motion [11–13]. Likewise, recent research on cohesive cell groups has revealed a rich spectrum of mechanisms driving collective migration – crucial in physiology (e.g., embryogenesis) and pathological processes (e.g., cancer metastasis). A few notable discoveries have been made in the collective cell motility field; one such discovery is contact inhibition of locomotion (CIL), whereby cells that come into contact with each other cease their migration towards their colliding partner before repolarizing and migrating away from each other [14, 15]. Another important discovery is contact following locomotion (CFL), whereby a cell actively follows the direction of its neighboring cell, essentially “trailing” behind it, so that the leader cell and the follower collectively migrate in a coordinated manner [16]. Lastly, a number of groups have reported on the supracellular organization of cell clusters or even pairs of cells [17, 18]. Despite these important findings, the emergent modes of collective migration have been less well-characterized compared to the migration patterns of individual cells [19]. How can we characterize the emergent types of collective migration and understand the molecular and physical basis that dictates which type of migration a cell collective will opt to adopt?

One of the simplest forms of collective migration can be observed in the amoeba *Dictyostelium discoideum* (Dd), which upon starvation initiates streaming migration, initially forming tandem pairs where one cell follows another, before progressing to chains of multiple cells, asters, and other structures [20, 21]. Dd serves as a widely studied model for amoeboid migration (also exemplified by neutrophils), characterized by the extension of a 3D pseudopod at the front of the migrating cell that drives rapid, dynamic directional movement [7, 22, 23]. In contrast, Madin-Darby canine kidney (MDCK) cells, an epithelial cell model, primarily undergo mesenchymal migration, which relies on the formation of 2D lamellipodia (flat, sheet-like protrusions) that typically facilitate slower, more adhesive forward movement [24–26]. Unlike mesenchymal migration, much less is known about the cell-cell or cell-matrix adhesion in migrating Dd, as Dd cells lack integrin-based adhesions typical of mammalian cells, and their cell-cell adhesion complexes are not as well characterized. Bastounis et al. showed that as Dd cells move cohesively in pairs in the direction of a chemical gradient [8] (Figs. 1(a)-(b)), with both cells in these pairs maintaining their autonomous single-cell signature 80% of the time. Autonomous cell migration is characterized by the formation of two diffuse cell-matrix adhesion sites, one at the front and one at the rear of the cell, allowing the cell to contract inwards (towards its center) [8]. Thus, two large but distinct adhesion sites are apparent at the front and rear of both cells of a migrating pair. Each cell acts as an individual contributor generating a characteristic traction force dipole, with a strong propulsion at the front and a strong drag at the rear (individual contributor (IC) mode; Fig. 1(b)). Still, the leader (i.e., front cell) and trailer (i.e., back cell) cell’s dynamics are not equal: the rear of the leader cell pulls the front of the trailer cell, dragging the trailer forward. Interestingly, adhesion sites are “recycled” so that the “old” adhesion site at the rear of the leader becomes the “new” adhesion site at the front of the trailer. The remaining 20% of the time, only two large adhesion sites remain in the pair: one at the front of the leader and another at the rear of the trailer (labeled S, Fig. 1(b)). In this state, the two neighboring traction force dipoles fuse into a single contractile dipole, with all propulsion in the leader and all resistance in the trailer. Thus, the pair in this state is in a supracellular motility configuration (S mode), with the leader and trailer cell behaving like the front and rear half of a single motile cell, respectively. Intriguingly, in the supracellular state, the pair’s speed decreases. In contrast to these observations in Dd, a recent study on the migration of chains of MDCK cells illustrates the opposite behavior [26]. That is, migration speed remains constant with increasing cell chain size. Moreover, nearly all the time, both MDCK cells in the pair exhibit supracellular motility (S mode), as evident by their traction force distributions (Fig. 1(c)).

**FIG. 1.**
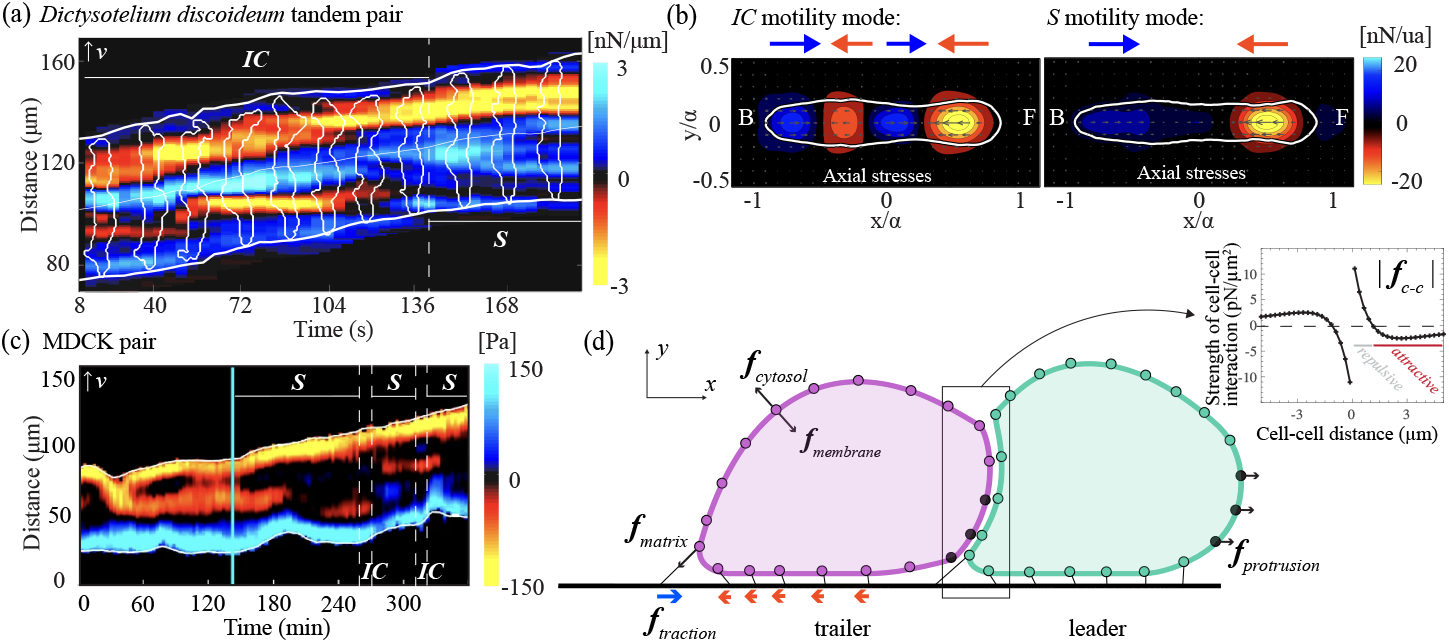
Traction stress dynamics of cohesive Dd and MDCK cell pairs moving on a flat surface based on *in vitro* experiments. (a) Representative kymograph of the axial traction tensions of a streaming pair of Dd cells chemotaxing on a soft 1.2 kPa matrix adapted from [8]. Blue color refers to tension facing towards the front of the migrating cell pair (positive value), and yellow color to tension facing towards the rear of the migrating cell pair (negative value). The white outlines mark the instantaneous position of the front, centroid, and rear of the doublet. The Dd doublet spends roughly 80% of its time moving in IC mode (two force dipoles), and the remaining time in S mode (single force dipole). (b) Mode-averaged axial traction stress maps (nN/unit area) in the pair-based reference frame for *N* = 14 Dd pairs moving in the IC mode (left) or S mode (right). The white contours show the average shape of the pair, and the front (F) and back (B) of the pair are also indicated. Experimental procedures and data analyses are in [8]. (c) Representative kymograph of the axial traction stresses in a pair of MDCK cells taken from [26]. The cyan line marks the beginning of the photoactivation of the Rho GTPase Rac1 in the front most cell (i.e., leader cell). The pair moves predominantly in S mode (single force dipole). (d) A side view schematic of the biophysical model of the migrating cell doublet, with polarization in a fixed direction (right). In the model, cells move according to the force balance in Eq. 1, which considers friction, the elastic response of the membrane structure, volume incompressibility, the actin-driven protrusion at the front, cell-matrix adhesion, and cell-cell adhesion. Black circles mark the protrusive region of each cell of the pair. The links underneath the cells represent cell-matrix adhesion bonds. The inset shows the magnitude of the cell-cell adhesion modeled as a Morse potential, characterized by a short-range repulsion and a long-range attraction

What physical mechanism is responsible for the different migratory modes adopted by Dd versus MDCK cell pairs? What is the role of cell-cell adhesion and cell-matrix adhesion in dictating the emergent collective cell migration mode of cell pairs? How is synchronization between cell-cell contact and cell-matrix adhesion achieved in cell chains? We attempted to construct a single model of the cell pair that captures the rich landscape of behaviors observed in a previous study with Dd pairs [8]. We used a mechanochemical modeling framework, originally built for modeling single cell Dd motility in [27], to describe our doublet’s motion and assumed that cell-cell adhesion exerts a combination of attractive (CFL-like) and repulsive (CIL-like) forces as others have done [14, 28]. Our model successfully reproduces both IC and S motility modes, demonstrating how variations in membrane mechanical properties and cell-matrix adhesion influence migration dynamics of cell pairs. Specifically, the model shows that coupling two “stepping” amoeboid cells (such as Dd) predominantly produces double dipole traction stress signatures, consistent with the emergence of the IC mode observed experimentally. On the contrary, coupling two mesenchymal cells (such as MDCK) favors a supracellular arrangement with a single traction force dipole, as in vitro. We further explored how additional perturbations in the model affect migration speed and the emergent motility modes. Our findings emphasize the significant role of cell-matrix adhesion in modulating migratory mechanisms, also in collective migration settings. Importantly, the model provides a framework to reconcile seemingly contradictory experimental observations, demonstrating the power of computational approaches in understanding complex cellular behaviors.

## II. MODEL

Each cell in the doublet is modeled using a biophysical framework of a cell polarized in a fixed direction, originally proposed in [27]. Because doublet migration was examined on a two-dimensional adhesive matrix in [8, 26], we restrict our framework to two dimensions and adopt a side view of the cell (Fig. 1(d)). The position of the *i*^th^ point on the membrane is denoted by ***X***^(*i*)^ = *x*^(*i*)^(*s, t*), *y*^(*i*)^(*s, t*), where *t* is time and *s* is the local parametric coordinate on the membrane structure. We use the convention that *i* =1 labels the front (leader) cell and *i* =2 labels the rear (trailer) cell. The discretization method used in simulation is summarized in SI Note A. The evolution of each cell boundary is described as an overdamped system, and we locally balance forces per unit area to write [29, 30]

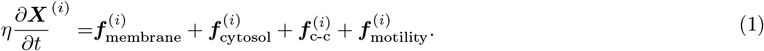

The left-hand side represents the friction force per unit area with viscous drag coefficient *η*. Below, we ignore the superscript distinction and describe the forces acting on a cell. The term ***f***_membrane_ models the cell boundary as a deformable structure, representing the membrane and cortical mesh, with an elastic response of stiffness *k* (units of force/length) and resting tension *γ* (units of force/length) [31–33],

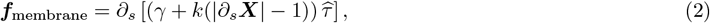

where 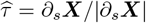 is a unit vector tangent to the cell boundary. The term ***f***_cytosol_ models cell incompressibility by penalizing deviations of cell area from its resting area *A*_0_ [27, 34],

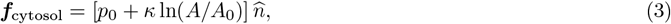

where *p*_0_ is given by

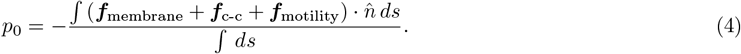

This force density points in the outward normal direction as indicated by the unit vector normal to the cell boundary 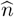. In the absence of forces due to protrusive activity, cell-matrix adhesion, and cell-cell adhesion, each cell equilibrates to a circle of area *A*_0_. The third term in the force balance is the cell-cell adhesion, which couples the cells in the doublet together. The last term represents the active motility force per unit area, which drives each cell front forward in a fixed direction, and incorporates the interaction between a cell and the adhesive matrix underneath.

### A. Cell-cell adhesion forces

Despite theoretical models of various types of interactions during collective migration [19, 28, 35, 36], it is unclear which interactions are required to capture the migration dynamics of small cell clusters. A recent study concluded that a number of cell-cell communication pathways result in an effective attractive-repulsive Morse potential [37]. The molecular underpinnings of this effective potential could stem from a combination of contact inhibition of locomotion [14] and contact following locomotion [16], enforcing an asymmetric distribution of Rho GTPases across the intercellular junction [38–40]. Rather than considering the detailed interactions of Rho GTPases across the cell-cell junction, we summarize cell-cell adhesion through a attractive-repulsive Morse potential similar to [37]:

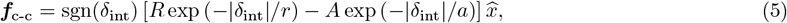

where *δ*_int_ is the horizontal distance between the neighboring cells (pointwise defined) and 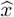 is a unit vector pointing in the horizontal direction. The first term inside the bracket describes repulsive forces, while the second term indicates attractive forces. The strengths of the forces are given by *R*, and *A*, while the spatial ranges for these forces are provided by *r*, and *a*, respectively. The cell-cell contact region is defined by the region of protrusive activity in the trailer cell and the corresponding region at the back of the leader cell, which minimizes the vertical separation to the trailer. Alternative cell-cell adhesion forces were considered in SI Note B (Fig. S1).

### B. Defining protrusions and cell-matrix adhesion forces

The motility force density incorporates protrusive activity due to the polymerization of F-actin filaments against the cell boundary [41, 42], balanced and transmitted through the cytoskeleton to adhesions sites, and cell-matrix adhesion,

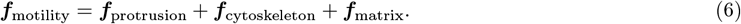

Migrating cells have a polarity, indicating the areas of the cell that are likely to protrude (“front-like”) and those likely to contract (“rear-like”). The molecular underpinnings can include asymmetric distribution of signaling proteins [5], reorganization of the cytoskeleton network [43], or a combination of the two. Rather than explicitly modeling the cell polarity dynamics [44, 45], we assume that a fixed region of the cell boundary, *P*, protrudes with a prescribed force-velocity relation due to the presence of branched F-actin filaments [41]:

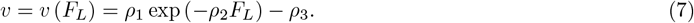

*F*_*L*_ is the force against the protrusion (load force) given by the average magnitude of forces in the protruding region. An equivalent way to formulate this relation is to assume the protrusive force is a function of the edge velocity; then, at the cell front, the force-balance equation in the direction of the motion becomes

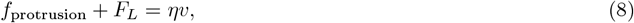

where 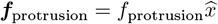. Lastly, we write the load force in one cell as

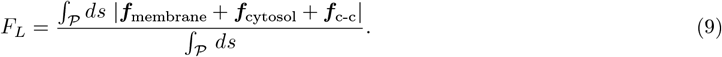

During pseudopod extension, the pseudopod must reach a minimal length of 10 *µ*m before attaching to the matrix. As presented, the protrusive forces are unbalanced and must be transmitted to the matrix underneath, in order to ensure conservation of momentum [46, 47]. We assume the protrusive forces are distributed uniformly to the region of cell-matrix contact, and mathematically write:

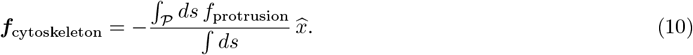

Finally, we assume each cell interacts with the underlying matrix through physical adhesion connections and a steric repulsive force to ensure the cell does not penetrate the matrix underneath. Hence,

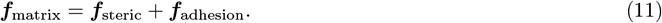

The steric repulsive force ensures the cell does not penetrate the matrix and is applied when the cell is below a certain threshold distance, ℒ_*s*_, in the vertical direction and is equal to

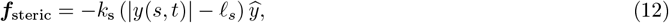

where *k*_*s*_ is the strength of the steric interaction and 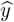 denotes the unit vector in the vertical direction.

In the experiments of [8], the cells crawl on a flat matrix coated with fibronectin. The authors of [27] showed that the apparent recycling of adhesion sites, evident in axial traction stress measurements, can only be explained by incorporating mechanosensitive cell-matrix adhesion. Similarly, we assume that these physical connections are force-sensitive and can weaken (“slip” bonds) or strengthen (“catch” bonds) with applied force [48]. This description can be written mathematically in different ways – here, we model the bonds as Hookean springs with force-sensitive dynamics, as in [27, 49]:

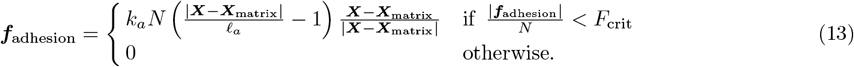

The adhesion force is written as a piecewise constant to capture the catch (bonds strengthen with applied force) and slip response (bonds rupture above threshold force), although other forms do not qualitatively change the emergence of “stepping” locomotion [27]. Here, *k*_*a*_ is the adhesion stiffness strength (units of force/area), ℒ_*a*_ is the rest length of the spring connecting the cell bottom to the matrix underneath, ***X***_matrix_ represents locations of adhesion bonds along the matrix, and *F*_crit_ is the threshold applied force density. When a bond forms, its position ***X***_matrix_ is set to be the location directly underneath the corresponding location of the cell boundary, thus exerting zero tangential stresses. For the lifetime of the bond, its position remains fixed along the matrix. The local bond density evolves dynamically following

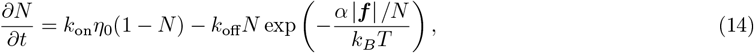

The first term represents the formation of new bonds at a rate proportional to a constant binding rate, *k*_on_ (units of M^*-*1^ time^*-*1^), and an unsaturated matrix ligand concentration, η_0_ (units of M). The second term represents force-dependent unbinding with a zero-force unbinding rate, *k*_off_ (units of time^*-*1^), and a microscopic length scale characterizing the unbinding transition, α. The total amount of force density applied to each adhesion site, ***f***, is proportional to the amount of stretch between the adhesion point on the cell and the ligand position on the matrix. We can then compute |***f*** | as |***f*** | = |***X*** *-* ***X***_matrix_| /ℒ_*a*_ *-* 1.

### C. Parameter setting

The Dd cell doublet simulations use baseline values listed in Table I; the values are chosen to reproduce stepping locomotion similar to amoeboid cells in [8, 27]. The friction coefficient *η* sets the crawling speed and is chosen to match the timescale of biological motion. The line tension *γ* is in good agreement with previous measurements of cortical tension [23, 50–52], and *k* is chosen so that deformation forces are of the same order of magnitude as tension forces. The constraint penalizing deviations from the preferred area, *κ*, is set relatively high to ensure the area of the default cell varies less than 1% from its target. As in our earlier work [27], the polymerization constants *ρ*_1_, *ρ*_2_, *ρ*_3_ are chosen to reproduce a force-velocity curve with a stall force per filament that corresponds to a range of 1 − 10 pN as observed in multiple studies [41, 42, 53, 54]. The strength of the steric cell-matrix interaction *k*_*s*_ is set to ensure the cell remains at a fixed horizontal distance ℒ_*s*_ from the matrix. Lastly, the adhesion cell-matrix parameters are calibrated to reproduce the stepping locomotion observed in singlet and doublets, as in [8, 23]. Measurements of exact intercellular forces across cell types are limited. We picked the CIL and CFL parameters in the Morse potential so that the maximum distance of the cell-cell contact region of migrating Dd doublets is approximately 0.5 *µ*m, which aligns with previous studies [55, 56]. The remaining parameters in Table I will be discussed in Section III as we consider their variations and effect on the doublet behavior.

**TABLE I.** Simulation parameters, along with the values that define the default for each cell in the doublet. The simulations are calibrated to behave similarly to Dd cells in [8, 23].

| Parameter | Value | Units |
| --- | --- | --- |
| $\eta$ | 72 | pN s/$\mu\text{m}^3$ |
| $\gamma$ | 800 | pN/$\mu\text{m}$ |
| $k$ | 800 | pN/$\mu\text{m}$ |
| $A_0$ | 179.55 | $\mu\text{m}^2$ |
| $\kappa$ | 800 | kPa |
| $R, A$ | 16, 4.8 | nN/$\mu\text{m}^2$ |
| $r, a$ | 0.75, 4 | $\mu\text{m}$ |
| $\rho_1$ | 11.5556 | $\mu\text{m}/\text{s}$ |
| $\rho_2$ | 0.0082 | $\mu\text{m}^2/\text{pN}$ |
| $\rho_3$ | 1.1111 | $\mu\text{m}/\text{s}$ |
| $k_s$ | 80 | kPa |
| $\ell_s$ | 0.6 | $\mu\text{m}$ |
| $\ell_a$ | 0.4 | $\mu\text{m}$ |
| $k_a$ | 800 | pN/$\mu\text{m}^2$ |
| $F_{\text{crit}}$ | 8400 | pN/$\mu\text{m}^2$ |
| $k_{\text{on}}\eta_0$ | 16.1074 | $\text{s}^{-1}$ |
| $k_{\text{off}}$ | 322.146 | $\text{s}^{-1}$ |
| $\alpha$ | $1.25 \times 10^{-4}$ | $\mu\text{m}^2/\text{pN}$ |
| $L_{\text{pseudopod}}$ | 10 | $\mu\text{m}$ |

### D. Traction stress measurements

We begin each simulation by initiating the cells near each other (i.e. 0.5 *µ*m apart), and numerically evolve the coupled equations of motion for 12 minutes. We allow the cells to relax for 0.3 minutes prior to applying protrusive forces for migration. To compare against the experiments, we track the spatiotemporal evolution of the cell length and traction stresses (force per unit area) for each cell. The traction stress is defined to be the opposite of the cell-matrix adhesion force density: ***f***_traction_ = *-****f***_matrix_, but only the tangential component of the traction stress is shown.

### E. Quantification of motility modes and related measurements

All simulations (singlet or doublet) are run for 12 minutes to ensure the migration of the cell doublet enters a steadystate dynamic. We calculated the velocities starting from 3 minutes to exclude a transient period from stationary to moving pairs.

In [27], we reported that a single moving cell exhibits a force dipole concentrated at both ends of the cell by exerting traction stresses on the underneath matrix. Traction stresses are plotted in a reference frame in which the coordinate axis *x* coincides with the major axis of the cell. In this coordinate system, in which the cell moves along the *x*-direction, the traction stresses are negative at the front of the cell and positive at the rear, consistent with a dipole of inward contraction. Similarly, we quantified migratory patterns by examining traction stresses of the doublet. As reported experimentally in [8, 26], we found that cell doublets can exhibit two distinct traction stress patterns: a single force dipole similar to a single moving cell called “S mode” (or supracellular mode), or two force dipoles arranged sequentially, called “IC mode” (or individual contributor mode). To classify the mode of migration in a Dd doublet, we adopt the following strategy. We found that the leader cell will demonstrate an inward contractile force dipole (i.e. 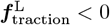 at the front and 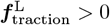 at the back) majority of the time. Moreover, the traction stress at the back of the trailer cell is always positive, and it is, therefore, sufficient to focus on the traction stress of the trailer cell to determine if the doublet is moving in mode S or mode IC. In particular, the migrating doublet is classified to be in:

1. IC mode: if traction stresses at the front of the trailer cell are negative;
2. If traction stresses are positive at the front of the trailer cell, then
  a. S mode: if traction stresses at all other locations are non-negative;
  b. IC mode: otherwise.

To quantify the motility modes of MDCK-like doublets, we used a modified scheme summarized in SI Note C. A freely available repository, containing C/C++ code used for simulations and MATLAB code used for motility mode quantification and visualization, can be found at https://github.com/yz-yingzhang/Collective_Migration_Cell_Chains.git.

## III. RESULTS

### A. The model predicts that a tandem migrating cell pair exhibits two distinct modes of migration

In the model proposed above, the single cell moves as a “stepping” cell, similar to what is observed during chemotaxis in amoebas crawling on a flat surface (Fig. 2(a) and S1 Movie) [7, 22, 57]. The cell migrates through a cyclical process, as reviewed in [22], which involves the extension of a protrusive front at the leading edge (termed pseudopod), adhesion of the polarized front to the matrix, whole cell body contraction, and rear retraction to push the cell forward. Simulations of a motile amoeba present notable oscillations in cell length over time, with a period and amplitude closely matching experimentally reported values [27] (Fig. 2(b)). The simulated singlet moves at a roughly constant speed of 11 *µ*m/min (the reported values for Dd are 9 − 12 *µ*m/min [22]). Also in agreement with experiments, traction stresses exerted on the matrix, and localized at the cell’s front and rear regions, are also closely tied to this motility cycle (Fig. 2(a)). Initially, these forces are directed inward and form a dipole, with negative stresses at the front and positive stresses at the back. During movement, distinct and recycled adhesion sites can be seen by the horizontal patches in the kymograph (white arrowhead, Fig. 2(c)). Another defining feature of this process is the dynamic turnover of traction adhesion sites: as the cell extends a new pseudopod, it establishes a new adhesion site at the front edge while retracting the rear by rupturing adhesions at the back (Fig. 2(a)). Subsequently, the adhesion sites at the former front edge of the cell are seemingly recycled to become the new rear traction adhesion sites, thus completing the cycle. The emergence of step-like locomotion depends on the force-dependent protrusive activity in pseudopod formation and the catch-slip nature of cell-matrix adhesion bonds.

**FIG. 2.**
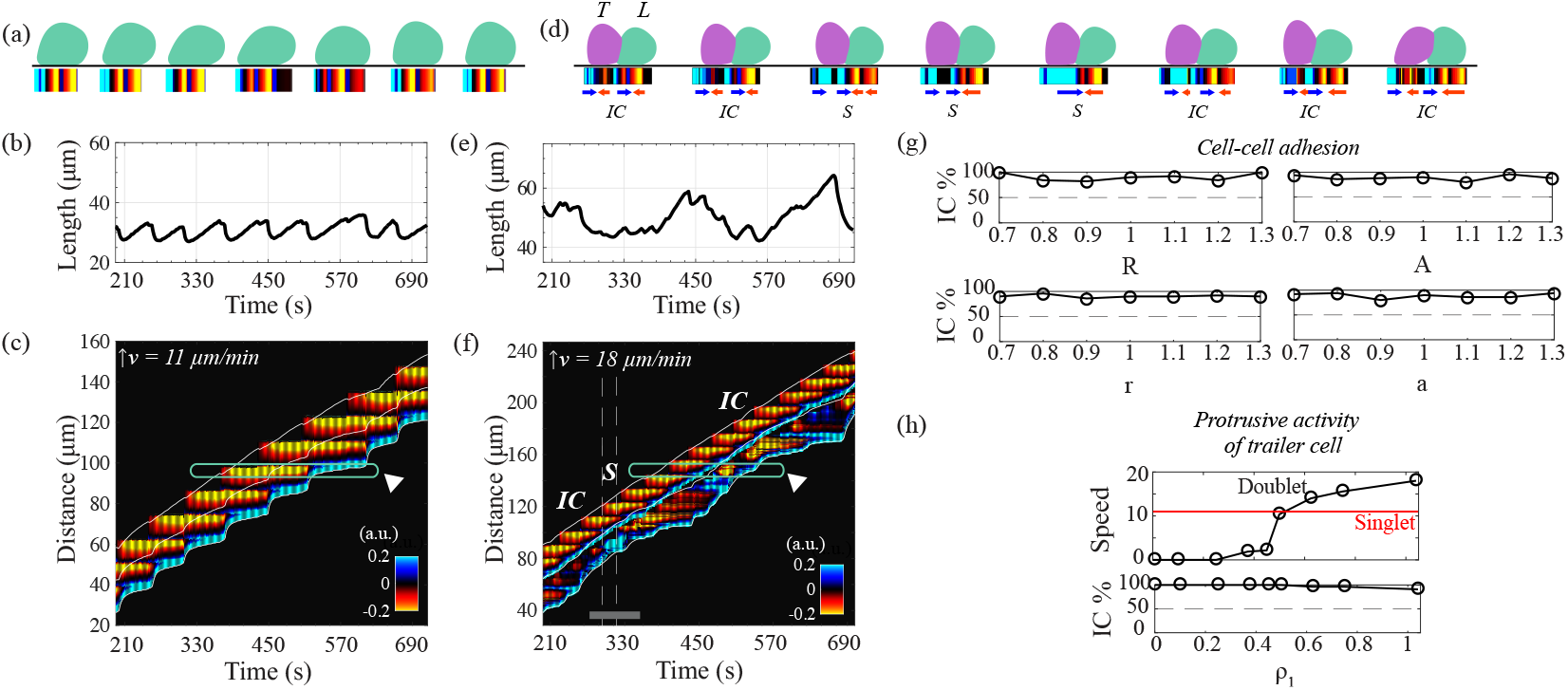
The model captures both IC and S migratory modes of cell pairs, with IC mode emerging the majority of the time (as observed experimentally), while cell-cell adhesion and protrusive activity of the trailer cell do not affect the mode distribution. Migratory patterns of a simulated single amoeboid cell ((a)-(c)) and doublet ((d)-(f)). Each case shows sample snapshots of cell morphologies and exerted axial traction stresses over a motility cycle ((a),(d)), line plots of length over time ((b),(e)), and kymographs of axial traction tensions ((c),(f)). The emergent motility mode is indicated by the text underneath the cell outlines, while the text above marks trailer (T) and leader (L) cell. White arrowheads in (c) and (f) highlight the appearance and disappearance of the distinct traction adhesion sites at a fixed location on the matrix. The gray bar in (f) identifies the region corresponding to cell outlines in (d). In both (c) and (f) the white solid inclined lines indicate the instantaneous position of the front, centroid, and rear of the cell chain. (g) Scatter plots of resulting percentage of IC mode when varying cell-cell adhesion forces by changing each of the four associated parameters, revealing that mode distribution does not depend on cell-cell adhesion. (h) Scatter plots of doublet speed and resulting percentage of IC mode with changes in the strength of protrusive forces in the trailer cell (T), modulated by parameter *ρ*_1_ as in Eq. 7. In (g) and (h), numbers on the *x* − axis label the fractions of parameter values relative to their baseline value listed in Table I.

But can our proposed model for a tandem pair simultaneously capture features of both the supracellular mode and the individual contributor motility mode, as observed for Dd cells in [8]? Using the baseline parameters for individual cells together with the cell-cell adhesion defined above, Figs. 2(d)-(f) plot one entire cycle of the doublet morphologies along with emergent pair length and traction stresses (also S2 Movie for a longer run). The doublet migrates persistently to the right while maintaining cohesiveness – the “leader” cell forms a well-defined protrusive pseudopod, while the “trailer” cell curves slightly outward into the leader cell, producing a convex cell-cell junction. In contrast to singlets, where oscillations in cell length are clearly visible (Fig. 2(b)), in doublets the oscillations are less well-defined (Fig. 2(e)). Fig. 2(f) shows the kymograph of the traction stresses over time along with outlines of the doublet front, center of mass, and rear. When the tandem pair migrates collectively, the doublet predominantly adopts an IC mode, where each cell behaves as an autonomous entity with a characteristic traction force dipole. Consistent with experimental observations, this mode of migration results in a double-dipole signature throughout most of the migration process (Fig. 2(f)). Distinct and recycled adhesion sites between the leader and trailer cell are seen in the kymograph. A fixed location on the matrix identified as an adhesion site in the leader cell is recycled to an adhesion site in the trailer cell (white arrowhead, Fig. 2(f)).

We quantified the migration mode as discussed in Section II E and found that the percentage distribution of IC (double traction dipoles) to S (unified traction dipole) is 95% − 5%. To illustrate the switch, we focus on a time right at the cusp of an IC-to-S transition (gray bar, Fig. 2(f)). The switch to S mode – a single traction force dipole – is induced when the trailer cell retracts due to strong pulling from the leader cell. The trailer cell recycles adhesion sites that are initially formed by the leader cell, highlighting a coordinated yet independent interaction between the two cells (white arrowhead, Fig. 2(f)). This finding is consistent with the recycling of distinct adhesion sites in doublets reported in [8].

The pair migrates faster (18 *µ*m/min) compared to singlets (11 *µ*m/min), a trend that deviates from experimental observations. To address this discrepancy, we systematically perturbed the cell-cell adhesion parameters by 30% (Fig. 2(g)). The emergent motility mode is largely unaffected by changes in the strength of attractive or repulsive intercellular contributions; the closest match to the experiments in [8] is 90% − 10%. We found that larger perturbations in any one cell-cell adhesion parameter result in stalled cells or numerical artifacts such as overlapping cells. Moreover, the doublet speed still remains largely unaffected (Figs. S2(a)-(b), SI Note D). Next, we aimed for a closer match to both the doublet speed and the reported 80% − 20% IC-to-S distribution. We thought we can reduce the doublet speed by lowering the strength of the protrusive force in one cell. Indeed, when the trailer cell’s protrusive activity is dampened with 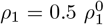, potentially due to a smaller pseudopod, this reduced the doublet’s speed (10 *µ*m/min). This result is in closer alignment with the experimental trend that Dd doublets move slower than individual cells [8]. While the doublet speed can be controlled by tuning the protrusive activity in the trailer, the emergent motility mode is largely unaffected by this parameter (Fig. 2(h)). While the protrusive activity can be controlled by ρ_1_, the strength of the protrusion can also be modulated by the area of the protrusive region. To examine the effect of varying the area of the protrusive region, we systematically changed its size by 30% (see SI Note E). We found that the motility mode remains largely unaffected by changes in the size of the protrusive region of both cells (Fig. S3, SI Note E). Taken together, our results suggest that the coupling of two stepping cells naturally gives rise to two motility modes, (1) an autonomous single-cell signature, the majority of the time, and (2) a supracellular signature indicative of mechanical fusing of the doublets, the remaining of the time.

Given the dynamic and largely unknown forces at the intercellular junction [19], we hypothesized that cell-cell adhesion may play a critical role in regulating the emergence of these modes. We investigated the effect of another cell-cell coupling scheme in regulating doublet migration (Fig. S1, SI Note B). We replaced the Morse potential with an alternative model that incorporates CIL, represented as a short-range steric repulsion, and CFL, represented as a long-range elastic-like attraction [28]. Our model results are qualitatively similar – the same two motility modes emerge in doublets. In contrast, when we investigated cells adhering to each other solely through linear elastic springs, the doublet struggles to maintain cohesiveness, often exhibiting separation or overlap. This occurs because linear elastic springs fail to provide sufficiently strong short-range repulsion to stabilize intercellular spacing. These findings suggest that the emergence of migration modes is robust to variations in cell-cell adhesion forces, provided both CIL and CFL are present, with CIL likely exerting a stronger influence than CFL.

### B. The strength of cell-matrix adhesion modulates the emergent motility mode both *in silico* and *in vitro*

Building on the evidence that cell-matrix interactions can influence the migratory mode in single cells [7, 27] (also S1 and S3 Movies), we hypothesized the same could be true in doublets. Fig. 3(a) quantifies the occurrence of IC mode while varying two cell-matrix adhesion parameters, bond formation rate (*k*_on_) and threshold rupture load (*F*_crit_), in both cells. Simulations revealed a clear trend: as either cell-matrix adhesion parameter, *k*_on_ or *F*_crit_, is increased, the doublets moved in IC mode more frequently, from 30% (filled star, Fig 3(a) and S4 Movie) to 100% (filled circle, Fig 3(a)). Increasing either cell-matrix adhesion parameter resulted in stronger cell-matrix adhesion bonds due to an increase in the equilibrium bond number. Conversely, lower values of *k*_on_ or *F*_crit_ reduced the cell-matrix adhesion strength, and led to a transition toward S mode.

**FIG. 3.**
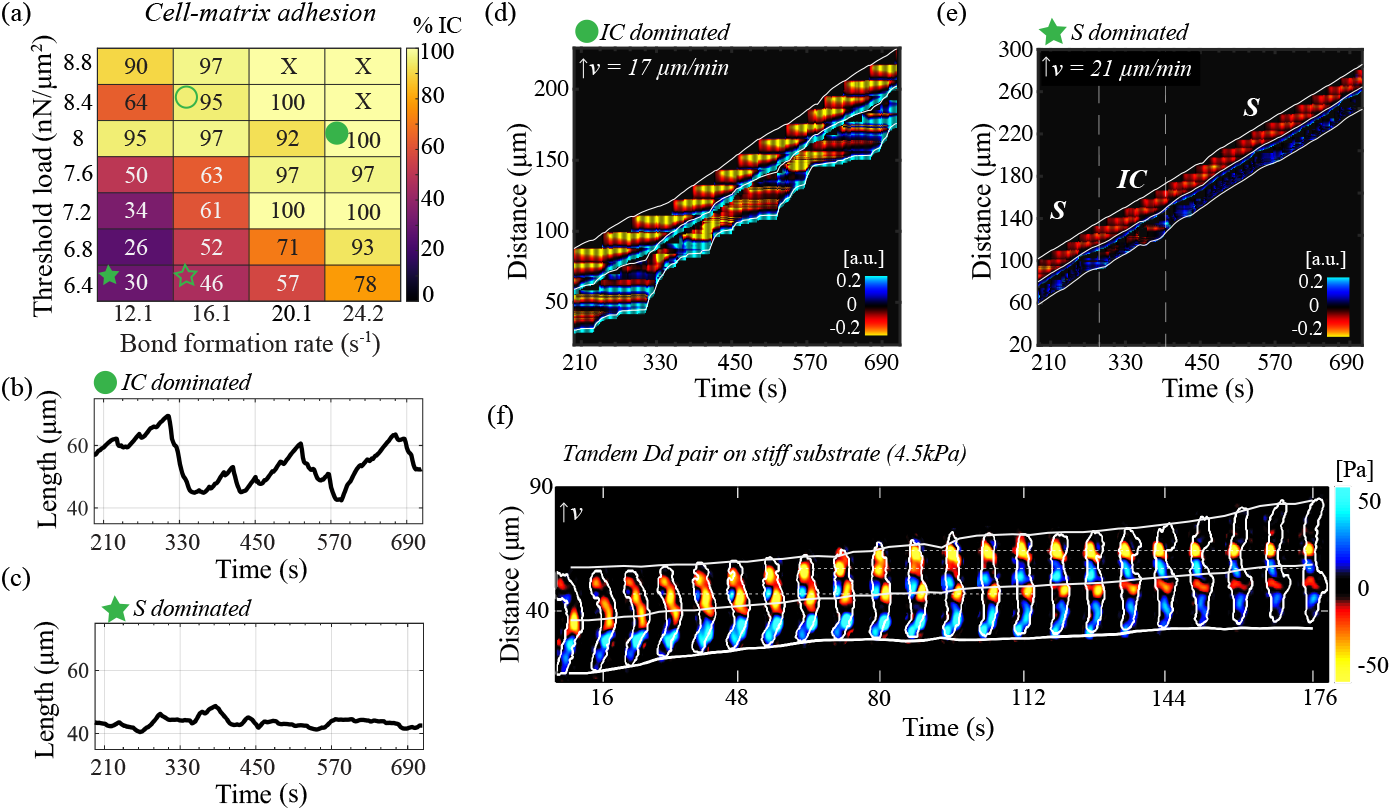
Increasing cell-matrix adhesion forces increased the prevalence of IC mode, both in the model and experiments. (a) Heatmap of IC mode occurrence (shown as percentage of total simulated time) with changes in the cell-matrix adhesion bond threshold load (*F*_crit_) and bond formation rate (*k*_on_). In (a), ‘X’ indicates stalled motility. The hollow circle represents cell-matrix parameters for a default stepping Dd doublet, while the hollow star marks a gliding Dd doublet. (b)-(c) Line plots of the two-cell chain length over time for two parameter choices, filled star and circle respectively in (a). (d)-(e) Kymographs of axial traction tension over time for the same two parameter choices as in (b) and (c). (f) Experimentally obtained kymograph of axial traction stresses (Pa) in a Dd doublet on stiffer 4.5 kPa substrate (where cells tend to increase traction stresses), demonstrating a predominant IC mode and large variations in cell length. In (d)-(f), white outlines indicate the instantaneous position of the front, centroid, and rear of the cell pair.

The effect of modulating adhesion strength is evident in both cell length dynamics and kymographs of the exerted axial traction stresses (Figs. 3(b)-(e)). Strong adhesion correlated with well-defined cell length oscillations (Fig. 3(b)) and strong traction dipole in each cell (Fig. 3(d)); these are defining features of IC mode. Weak adhesion, by contrast, resulted in diminished amplitude in cell length oscillations (Fig. 3(c)) and a shift toward coordinated traction dynamics of the S mode. We note that doublets in the S mode also exerted weaker traction against the matrix, and the recycling of traction adhesion sites was not observed (Fig. 3(e)). These findings align closely with the experimental results in Fig. 3(f); tandem Dd doublets migrating on stiffer substrates were more likely to exhibit IC mode. This agreement between model simulation results and experiments underscores the importance of cell-matrix adhesion strength as a determinant of collective motility mode. Strong cell-matrix adhesion enables cells to maintain independent traction stresses, reinforcing the autonomous dynamics required for IC mode, while weaker adhesion promotes a coordinated, supracellular behavior.

### C. The mechanical properties of the cell membrane and cortex modulate the predominant migratory mode

Given that we found that cell-matrix adhesion strongly influences the predominant collective migratory mode, next, we focused on the role of intrinsic cellular mechanical properties. We expected that varying cell membrane rigidity – γ and *k* parameters together – should affect the doublet’s migration, since these parameters directly correlate with how strongly cells pull (push) on their environment. For example, when γ, and *k* are low, i.e. the cells are soft, deformations are not very costly and can occur more easily, leading to rupturing cell-matrix adhesion bonds with ease. We expected stiffer cells to have a harder time to deform, and thus struggle to pull and rupture adhesion bonds; stiff cells should more likely “behave” similarly to doublets with stronger adhesion bonds and thus engage in individual contributor arrangement (IC mode). To test this hypothesis, we first perturbed our default Dd doublet (hollow circle, Fig. 3(a)), by changing γ, and *k* simultaneously by 10 and 20%, respectively, and then simulated its migration. The circles in Fig. 4(a) present the percentage of time spent in IC mode with variable membrane rigidity – there was essentially no effect on migration mode because the doublet is in a strong cell-matrix adhesion regime that dominates the effect of changing membrane properties. In a second perturbation, both cells in the doublet have a lower cell-matrix bond rupture threshold, 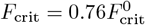 (hollow star, Fig. 3(a)), resulting in the doublet moving with 46% − 54% IC-to-S distribution (diamonds, Fig. 4(a)). In agreement with our hypothesis, we observed that soft pairs prefer S mode, while stiff pairs preferred IC mode. This finding led us to conclude that stiff cells mirror the behavior of cells with a higher rupture threshold – meaning that deformations were more costly, the adhesion bonds were pulled on less strongly and were less likely to rupture. Our findings demonstrate that altering intrinsic cellular properties, such as membrane elastic stiffness and tension, could significantly impact the emergence of migratory mode. Specifically, softer cells were more likely to move as a supracellular unit. These observations highlight that changes in membrane mechanics, which impact the ability of cells to pull on the matrix underneath, are analogous to variations in cell-matrix adhesion bonds. This reinforces our earlier conclusion that, through direct or indirect manipulations, the dynamic interplay between cellular stiffness and cell-matrix adhesion stands out as the central regulator of the migration of small groups.

**FIG. 4.**
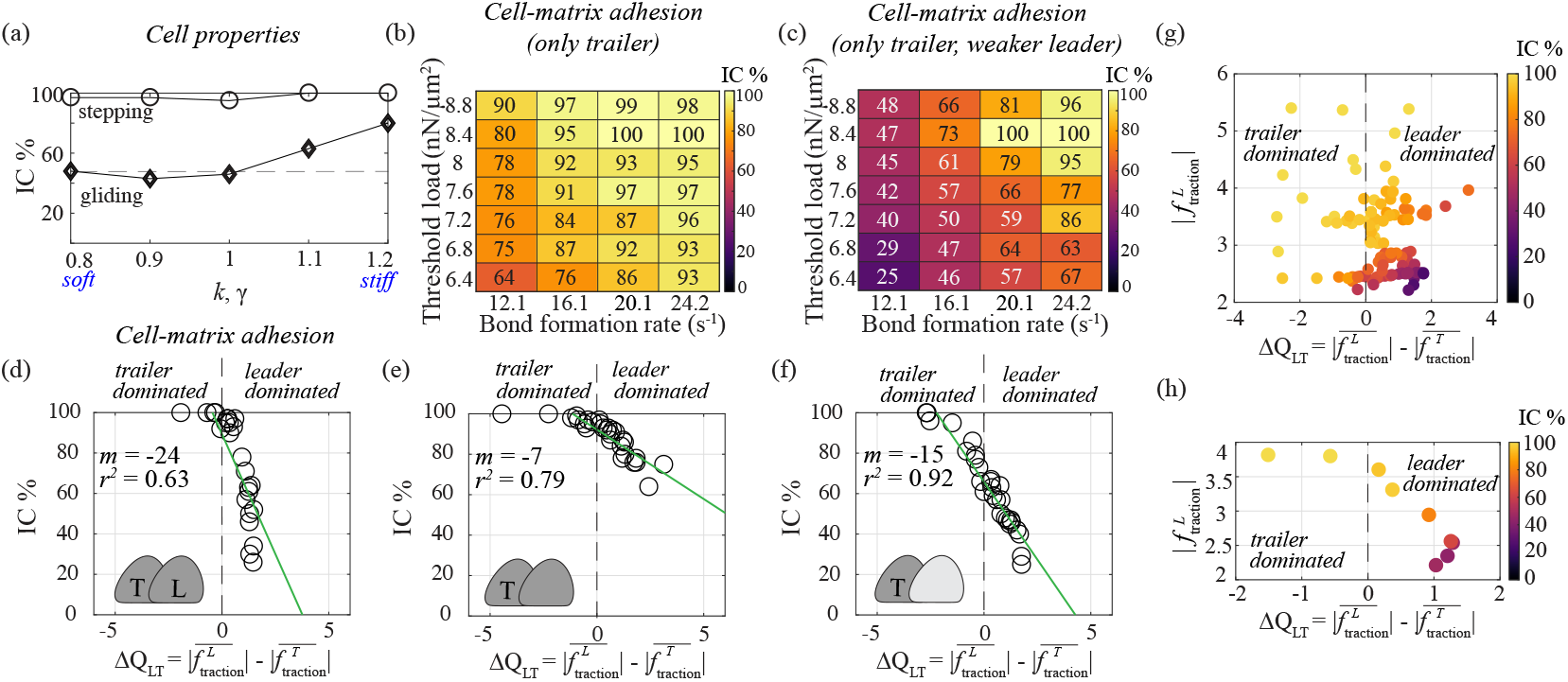
*In silico* studies revealed that weakly adherent cell pairs with a “pulling” leader produce supracellular arrangement (S mode). (a) Scatter plot shows the distribution of simulated Dd motility mode for varying membrane rigidity. Both tension *γ* and elastic stiffness *k* are simultaneously varied from their baseline values in default doublet (circles) and doublet weakly adhering to the matrix (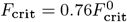, diamonds). Numbers on the *x −* axis label the fractions relative to baseline values in Table I. (b)-(c) Heatmaps of IC mode occurrence (shown as percentage) with changes in cell-matrix adhesion in: only trailer cell (b), and only trailer cell but with a leader cell with reduced threshold load of cell-matrix adhesion bonds 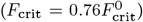 (c). (d)-(f) Scatter plots of occurrence of IC mode as a function of the difference in time-averaged traction stresses between the leader and trailer cells for cases in Fig. 3(a), Figs. 4(b), and (c), respectively. Best-fit lines are shown for the simulated data with slope and goodness-of-fit statistics (green lines). (g) Scatter plot of time-averaged leader traction stresses with varying cell-matrix adhesion forces represented as a function of traction stress difference, corresponding to data shown in (d)-(f). (h) Scatter plot of time-averaged leader traction stresses as a function of traction stress difference with varying membrane rigidity, corresponding to data shown in (a). In (g) and (h) marker face color represents percentage of IC mode.

### D. A pusher-puller mechanism between the leader and trailer cells of a pair underlies the motility mode distribution

Next, to uncover how individual cells contribute to the emergent locomotion phenotype, we systematically examined variations in cell-matrix adhesion and their effects on motility modes. Starting with a default leader cell coupled to a trailer cell with varying cell-matrix adhesion parameters, we observed that weakened adhesion in the trailer cell reduced the prevalence of IC mode. However, in most cases, the cells still acted as individual contributors, as indicated by the double traction force dipole (Fig. 4(b)). This observation led us to hypothesize that further reducing the leader cell’s adhesion strength would amplify S mode occurrence. Consistent with this hypothesis, when the leader cell’s interaction with the matrix was reduced 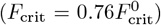, S mode dominated, as evident by a significant decrease in IC mode occurrence (Fig. 4(c)).

To understand the mechanism, we calculated the time-averaged axial traction stresses for each cell in the doublet over 200–720 seconds and defined the difference between these forces as the “traction stress difference”

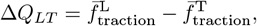

where *f*_traction_ is the axial component of the traction stress vector, ***f***_traction_. A positive Δ*Q*_*LT*_ indicates a stronger “pulling” leader cell relative to the trailer cell. Across all conditions – variations in adhesion parameters for both cells, the trailer cell only, or the trailer cell coupled with a weaker-adhering leader cell – we quantified the occurrence of IC mode as a function of 6Δ*Q*_*LT*_ (Figs. 4(d)-(f)). The results revealed a clear pattern: IC mode dominated when the trailer cell exerted stronger traction stresses, indicating a “pushing role”, while S mode emerged when the leader cell assumed a “pulling role” with weaker adhesion strength, yet stronger than the trailer cell. The negative slope of the best-fit line highlights the increased likelihood of supracellular organization (S mode) with a strong leader cell, relative to the trailer cell. Furthermore, Fig. 4(g) shows that the absolute traction stress magnitude of the leader critically determines the prevalence of S mode. In our model, supracellular migration emerged when the leader cell weakly adhered to the substrate but exerted higher traction than the trailer cell, resembling a “front-drive” pulling mechanism.

We extended this analysis to variations in cell membrane properties, examining their effects on motility modes. We found that the switch from IC-to-S mode remains governed by the pusher-puller dynamic (Fig. 4(h)). IC mode dominated when the trailer cell acts as a strong pusher, while S mode emerged when the leader cell assumes a pulling role. Taken together, our results demonstrated that a pusher-puller mechanism governs the emergent motility modes. Specifically, a contributing pushing trailer cell promoted IC mode, while a weakly adhering leader cell and a weaker trailer cell facilitated S mode. These findings underscore the central role of leader-trailer traction stress dynamics in regulating collective migration patterns.

### E. The model reproduces the traction stress dynamics of both Dd and MDCK-like migrating pairs

The simulated Dd tandem pair predominantly migrated in IC mode, characterized by two traction force dipoles, in agreement with the experimental findings of Dd streaming pairs in [8]. In [26], MDCK cell doublets appear to favor S mode, with a mode distribution estimated by us to be 95% − &5% IC-to-S (analysis in Section II E applied to Fig. 1(c)). Why do MDCK doublets exhibit a different distribution in migratory patterns? A few reasons could include that these cells move in a more mesenchymal fashion with well-defined adhesion complexes to adhere to neighboring cells and the matrix underneath, and do not form pseudopods, but rather use broad, flat lamellipodia confined to the very tip of the polarized cell [24, 25]. To mimic this, a few changes were made to our model: (1) the criterion for pseudopod length was removed, (2) protrusive activity was further spatially restricted to a region defined by *y* ≤ 3 *µ*m at the cell front, (3) adhesion stiffness strength, *k*_*a*_, was doubled at the cell front of the leader cell to replicate strong adhesion bonds near the lamellipodium. With these modifications, the migration pattern of an individual cell shifted from the stepping-like dynamics to a distinct gliding mode, characterized by the absence of oscillations in cell length (Figs. 5(a) and S5 Movie). Moreover, the kymograph of traction stresses over time was marked by the absence of a distinct, persistent traction adhesion site at the front (Fig. 5(b)), and the lack of recycling of these sites.

**FIG. 5.**
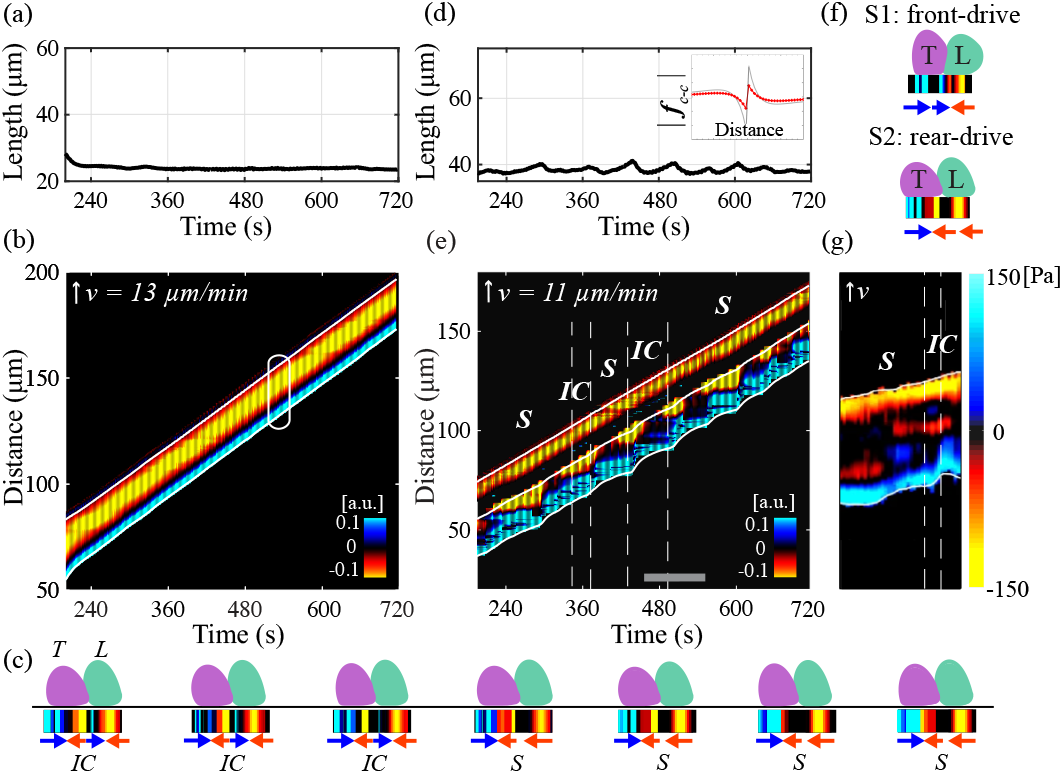
Simulated MDCK-like motile cell pairs utilized the supracellular arrangement (S mode) more frequently, but with a rear-drive, rather than a front-drive, mechanism. Migration patterns of a single MDCK-like cell ((a)-(b)) and a doublet ((c)-(e)). Each case shows plots of chain length over time ((a),(d)), and kymographs of axial traction stresses over time ((b),(e)). White oval is a hypothetical single cell outline. Inset in (d) shows the modified Morse potential (red) compared to default (gray). White outlines in (b) and (e) are the instantaneous position of front, centroid, and rear of the cell chain. Gray bar on horizontal axis in (e) labels the region corresponding to pair outlines in (c). In (c), in addition to the cell outlines and traction stresses, the emergent motility mode is indicated by the text underneath the cell outlines and the text above marks trailer (T) and leader (L) cell. (f) Sample morphology and traction stresses of the two types of supracellular modes: S1 (front-drive) and S2 (rear-drive). (g) Selected portion of MDCK kymograph data in Fig. 1(c).

To extend our model to doublets (Fig. 5(c)), a few additional changes were made in coupling the cells together (SI Table I, SI Note F). While the cell-cell contact region has roughly the same spatial spread as for amoeba pairs, it includes both protrusive and non-protrusive locations on the membrane since the polymerization region is smaller than in the case of amoeboid cells. The cell-cell adhesion parameters were adjusted to ensure the cohesiveness of the group by reducing both the CFL and CIL strengths (inset, Fig. 5(d)). Importantly, we found that to ensure the cells do not crawl on top of each other, we needed to not allow cell-matrix bond formation at the rear of the leader cell in the region of intercellular contact, i.e. where it links to its neighbor.

We repeated our analysis and quantified the emergent migratory patterns of the simulated MDCK-like doublets and observed the presence of two migratory modes based on patterns of traction stresses (Figs. 5(c)-(e) and S6 Movie). Overall, MDCK-like doublets migrated more slowly than individual singlets (11 *µ*m/min versus 13 *µ*m/min). With the parameters discussed above, S mode occurred 80% of the time and IC the remaining time (Figs. 5(c) and (e)). The doublet length continued to show low-amplitude oscillations (Fig. 5(d)), dissimilar from what was observed in Dd pairs. Notably, the supracellular organization is of a different flavor than that exhibited by the amoeba pair (S2 in Fig. 5(f)). The traction pattern, “S2”, is characterized by a force dipole in the trailer cell and a single traction site for the leader – referred to as a “pusher” or rear-drive doublet. This is different from the amoeba S mode, referred to as “S1” onward, which is a “puller” or front-drive doublet with a force dipole in the leader cell (S1 in Fig. 5(f)). The outlines of the doublet front and rear instantaneous position show an interesting feature – the trailer cell exhibits a stair-like pattern at its rear, reminiscent of “stepping” dynamics (Fig. 5(e)). This feature was also displayed in the *in vitro* MDCK doublet migration assay of [26] (Fig. 5(g)). While in some instances the traction stresses suggested a rear-drive mechanism under Rac1 GTPase optogenetic activation in [26], we could not confirm this conclusively without further analysis of the experimental data. Given the different migratory patterns of these MDCK-like cells compared to amoebas *in silico*, we wanted to probe the effect of parameter variations on migration and ascertain whether the pusher-puller analysis can provide additional mechanistic insight.

### F. The supracellular mode dominates when the trailer cell is a strong “pusher” for MDCK-like pairs

To explore the impact of cell-matrix adhesion, we adjusted the adhesion parameters for both cells simultaneously. Similar to the results for amoeba pairs, simulations of variations in cell-matrix adhesion parameters in MDCK-like doublets revealed distinct preferences for migration modes. Specifically, the S mode is favored when cell-matrix bonds exhibit a high threshold load but form rapidly, while the IC mode predominates under conditions characterized by a low threshold load and a slow bond formation rate (Fig. 6(a)). Examining the percentage of IC mode in relation to the difference in time-averaged traction stresses between the cells, Δ*Q*_*LT*_, we found little evidence showing any dependency between the mode and the pusher-puller dynamics (Fig. S4(a), SI Note G). To understand this ba?ing result, we simulated two additional scenarios – a default leader cell coupled to a trailer cell with changing cell-matrix adhesion parameters and a default trailer cell coupled to a leader cell with changing cell-matrix adhesion parameters. When cell-matrix adhesion parameters varied only in the trailer cell, we found that migratory modes were largely unaffected, with IC mode dominating (Fig. S4(b), SI Note G). However, changing cell-matrix adhesion parameters in the leader cell promoted the emergence of S mode (Fig. S4(c), SI Note G).

**FIG. 6.**
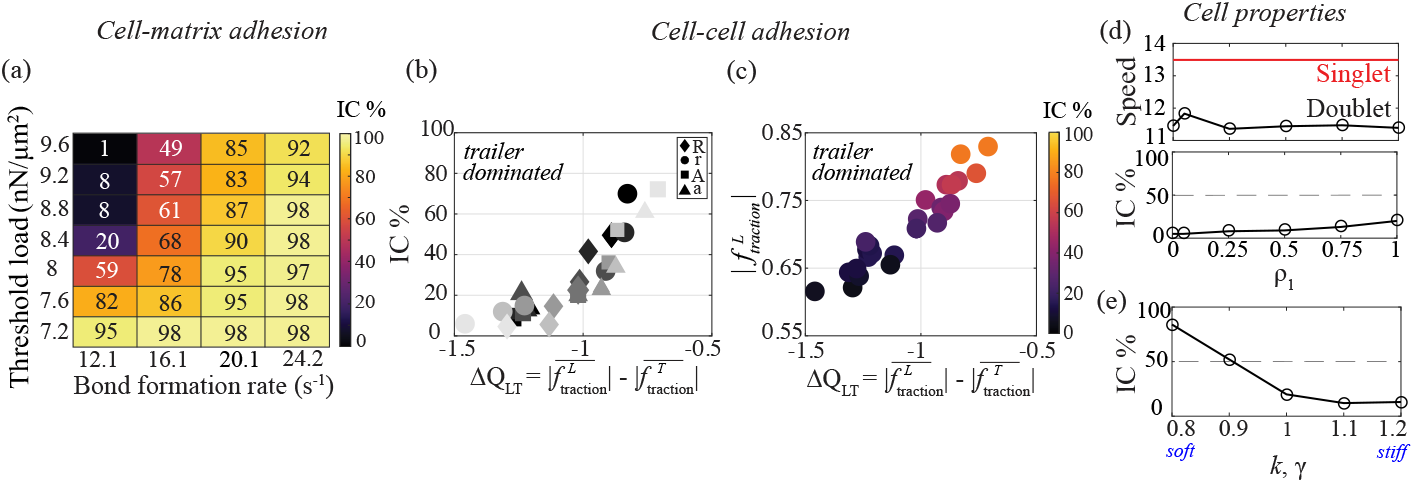
Unlike the Dd cell pair, the distribution of motility modes in MDCK-like doublets showed sensitivity to changes in cell-cell and cell-matrix adhesion forces *in silico*. (a) Heatmap of quantification of IC mode occurrence with changes in cell-matrix adhesion parameters. (b) Scatter plot of IC mode percentage as a function of traction stress difference with changes in cell-cell adhesion strength. Each symbol marks one change in one parameter in the cell-cell adhesion force. Darker color is a parameter increase by 10 or 20%, while lighter color is a parameter decrease by 10 or 20%. (c) Scatter plot of leader’s time-averaged traction stress as a function of the traction stress difference for simulations with varying strength in cell-cell adhesion. The color indicates the percentage of IC mode (color bar). (d) Plots of doublet speed (compared to singlet speed) and resulting IC mode percentage as a function of protrusive activity (*ρ*_1_) in the trailer. (e) Plot of IC mode percentage as a function of the cell membrane rigidity in both cells. In (d) and (e), numbers on the *x* − axis label the fractions of parameter values relative to their baseline values listed in SI Table I.

Although we observed a negative dependence between percentage of IC mode and the time-averaged traction stress difference as the trailer cell’s cell-matrix adhesion was varied, the changes were minimal (Fig. S4(d), SI Note G). Interestingly, when the leader’s cell-matrix adhesion was altered, we observed a positive dependence between the percentage of IC mode and the time-averaged traction stress (Fig. S4(e), SI Note G). These results suggest that cell-matrix adhesion of the leader cell might play a more prominent – but not dominant – role than the trailer cell in the emerging motility modes.

Strikingly, and unlike our prior findings in the amoeba pairs, changing the cell-cell adhesion parameters resulted in a redistribution of migration modes (Fig. S4(g), SI Note G). A reduction in CIL strength (*R*) or CIL spatial range (*r*) led to an increased prevalence of S mode. Conversely, an increase in CFL parameters (*A, a*) promoted IC mode. These changes in cell-cell adhesion had no statistically significant impact on doublet speed (Fig. S2(c), SI Note D). To further investigate these dynamics, we examined the time-averaged traction stress differences between the leader and trailer cells (Fig. 6(b)). A decrease in CIL parameters or an increase in CFL parameters enhanced the trailer cell’s role as a pusher, as indicated by Δ*Q*_*LT*_ < 0. Notably, the prevalence of S mode increased as the trailer cell exerted greater pushing forces relative to the leader cell (Fig. 6(c)), a trend that contrasts with the observations in Dd pairs. This finding reveals a reverse dependence on pusher-puller dynamics between Dd and MDCK-like doublets, highlighting their distinct migratory strategies.

Finally, we examined the influence of intrinsic cellular properties on migration dynamics in MDCK-like doublets. We analyzed the effects of changing the trailer cell’s protrusive activity (Fig. 6(d)). While an increase in the trailer cell’s protrusive forces did not significantly affect the overall doublet speed, it did introduce a slight bias toward IC mode. This finding suggests that more actively protrusive trailer cells induce the leader to adopt a more autonomous migration pattern. In this regime, the prevalence of S mode increases as the trailer cell’s pushing force intensifies, mirroring the effects observed with variations in cell-cell adhesion. Conversely, when the trailer cell’s polymerization strength falls below 25%, its ability to maintain autonomous movement is compromised, necessitating reliance on the leader cell’s pulling forces to sustain motility. This dependence leads to a higher frequency of supracellular (S) mode. Analysis of cell membrane rigidity variations revealed yet another fundamental difference between MDCK-like and amoeboid doublets. Specifically, in contrast to Dd cells (Fig. 4(a)), MDCK-like doublets composed of softer cells predominantly exhibited IC mode, whereas stiffer cells favor S mode (Fig. 6(e)).

### G. For longer cell chains, IC motility mode prevails for both *in silico* and *in vitro* Dd pairs

Having confirmed our model’s ability to replicate experimental findings for Dd doublets [8], we extended our investigation to longer cell trains. Using Dd cells with the baseline parameters listed in Table I, simulations revealed that the group often moved persistently to the right, but the rear-most cell detached from the matrix. This detachment was attributed to ineffective transmission of intracellular forces, as the inner cells acted as blockages due to equal regions of coupling at their front and back. To address this issue, we introduced a sequential 10-second delay in the protrusive activity of successive trailers. This modification ensured effective force transmission in the intercellular region, resulting in cohesive migration with all cells remaining attached to the matrix underneath (Fig. 7(a), S7 Movie).

**FIG. 7.**
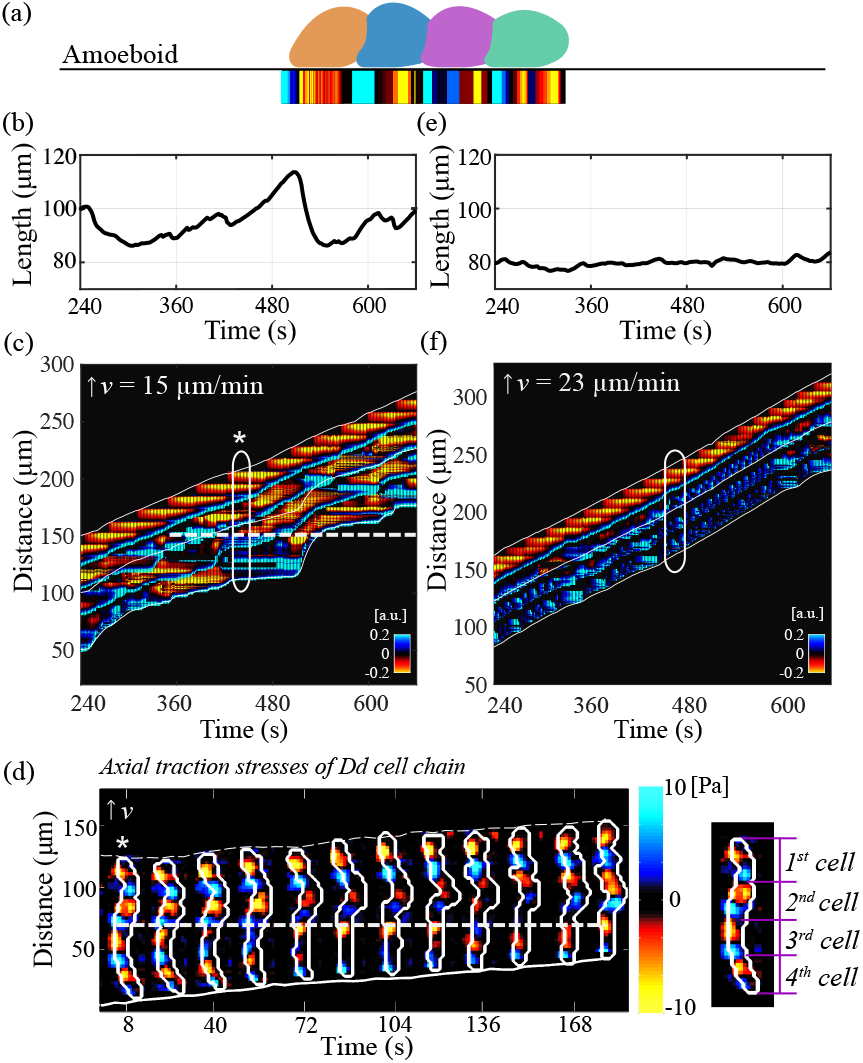
Both *in silico* and *in vitro* experiments with Dd showed that longer cell chains (e.g. 4-cell chain) move using multiple force dipoles and exhibit recycling of distinct, stationary adhesion sites. (a) Snapshot of morphologies and exerted traction stresses of a four- (amoeboid) cell chain. The migration patterns are shown in the default parameter case ((b)-(c)) as well as one with weaker cell-matrix adhesion (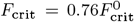, (e)-(f)). (b),(e) Line plots of total chain length over time. (c),(f) Kymographs of axial traction stresses over time. The white ovals show a hypothetical chain outline. (d) Representative *in vitro* experiment of the migratory patterns of Dd 4-cell chain showing axial traction stresses (Pa) exerted on a 1.2 kPa substrate. The inset on the right shows the four, distinct traction dipoles at the first instance of time shown on the kymograph (see asterisk). White outlines on the kymograph in (c) and (f) indicate the instantaneous position of the front, centroid, and rear of the cell chain. Dashed, horizontal, white lines in (c) and (d) mark a certain diffuse traction adhesion site which initially was the adhesion traction site of the second cell in the chain and eventually became an adhesion site of the fourth cell, and throughout the experiment remained at a fixed location on the matrix.

Using the same cell-matrix adhesion parameters as in the doublet simulations – strong adhesion (Fig. 2(f)) and weak adhesion (Fig. 3(e)) – we observed distinct behaviors. Compared to doublets, the length of the 4-cell chain exhibited fewer oscillations with longer periods and larger amplitudes (Fig. 7(b)). Each cell generated its own distinct traction force dipole, producing a kymograph pattern of four dipoles (Fig. 7(c)). Adhesion recycling was also evident in longer cell chains: a traction adhesion site initially associated with the second cell eventually became an adhesion site for the fourth cell (white dashed line, Fig. 7(c)). Occasionally, fusion of traction stresses between two neighboring cells resulted in three instead of four dipoles (asterisk, Fig. 7(c)). These motility patterns predominantly aligned with IC mode. The leader cell consistently exhibited a strong, persistent traction dipole, and these observations – distinct traction force dipoles, adhesion site recycling, and predominant IC mode – matched *in vitro* findings for Dd 4-cell chains (Fig. 7(d)). Consistent with our *in silico* findings in doublets, weakening cell-matrix adhesion increased the prevalence of S mode in 4-cell chains. Simulations with reduced adhesion strength 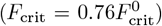 demonstrated a shift to S mode, characterized by trailer cells generating only positive traction stresses and reduced oscillation amplitudes in cell chain length (Figs. 7(e) and (f), and S8 Movie). The leader cell maintained a persistent traction force dipole, emphasizing its role as the “puller” in supracellular migration. These results suggest that the underlying mechanisms governing Dd motility modes are conserved across different cell train lengths.

We also extended this analysis to MDCK-like 4-cell chains (Fig. S5(a), SI Note H, S9 Movie). In default adhesion conditions, these chains exhibited small amplitude oscillations in cell chain length and migrated at speeds similar to singlets (Fig. S5(b), SI Note H). The corresponding kymograph revealed multiple dipoles, with S mode observed in the first two cells and IC mode in the remaining cells (Fig. S5(c), SI Note H). When cell-matrix adhesion was strengthened to mimic conditions with 1% IC prevalence in doublet simulations (Fig. 6(a)), the chains shifted further toward S mode with more pronounced cell chain length oscillations (Figs. S5(d)-(e), SI Note H). In agreement with findings in MCDK-like doublets, strong cell-matrix adhesion favors S mode in longer MDCK-like cell chains.

## IV. DISCUSSION

Bastounis et al. revealed the distinct migratory dynamics of streaming amoeboid Dd cells crawling on a flat elastic matrix by the dynamics of the distribution of axial traction stresses that these cells exert as they migrate. Here, we have proposed a simple biophysical model of the cell pair and showcased its successful recapitulation of observations in [8]. Our model correctly reproduces the recycling of distinct traction adhesion sites and the appearance of two distinct traction adhesion patterns: the S motility mode, characterized by a single supracellular force dipole, and the IC motility mode, which is illustrated by two force dipoles, one for each cell of the pair. We found that although cell-cell adhesion has little effect on which motility mode dominates, varying cell-matrix adhesion parameters does modulate the dominant motility modes. Increasing cell-matrix adhesion forces, specifically slow-forming adhesion bonds with a high threshold rupture load, results in pairs that move in IC mode, rather than S, over 90% of the time. Similar to these *in silico* findings, *in vitro* Dd pairs placed on a stiffer matrix, where cells tend to increase traction stresses [8], move in IC mode predominantly. The same model, with modifications to the protrusion and intercellular region assumptions, can also recapitulate the migration patterns of MDCK-like cells which, unlike Dd, undergo mesenchymal motility and are typically slower and more adhering to their matrix [26]. Our model predicts that these cell doublets move more frequently than amoeboid cells in S mode, similar to what was reported by [26] in MDCK chains upon localized Rac1 GTPase activation to induce motility.

The behavior of Dd and MDCK-like doublets is radically different, with cells switching from largely IC mode for Dd to more S mode for MDCK (Figs. 2(d) and 5(c)). We have been able to recapitulate the MDCK-like doublet dynamics with very minimal assumptions, including a thin protrusion region and local inhibition of cell-matrix adhesion in the intercellular region. The latter assumption seems well-supported by existing literature showing that E-cadherin engagement inhibits fibronectin adhesion [58]. In fact, a careful interplay between cell-matrix and cell-cell adhesions to maintain cohesive collective cell migration has been suggested by others [59, 60]. Our model suggests that this simple assumption has important ramifications on the emergent motility mode of migrating cell groups. In Dd, our analysis shows that the S mode is preferred when leader cells exert low traction stresses, yet stronger than those exerted by trailer cells. We refer to this S mode as a front-drive mechanism, with a force dipole in the leader cell and a positive traction adhesion site in the trailer cell (Figs. 2(f) and 5(f)). This is the opposite of what we observe in the S mode arrangement of MDCK-like cells (Figs. 5(e) and (f)), where the force dipole appears with the trailer cell. We refer to this mode as a rear-drive mechanism. Lastly, congruent with our findings in amoebas [8], we found that increasing the bias in the rear-drive mechanism, i.e. trailer cells that exert stronger traction adhesion forces than leaders, results in a preference for the S mode. Thus, our model proposes a unified framework that highlights the differences and similarities between migration strategies for cohesive cell pairs crawling on flat matrices.

MDCK and Dd cell types are very different cell types. MDCK cells exhibit strong cell-cell adhesion, mediated by well-characterized adhesion protein complexes. In contrast, much less is known about the adhesion mechanisms in Dd. Notably, Dd lacks integrin-based adhesions, a hallmark of mammalian cell-matrix interactions, and its precise mechanism of surface attachment remains unclear. Additionally, the way Dd cells adhere to each other is not as well studied as are the cell-cell junction complexes of mammalian epithelial cells, like MDCK. Similarly, the pseudopod membrane extension of amoeboid cells is also poorly characterized, in contrast to the sheet-like, flat, membrane extension – the lamellipodium – of MDCK cells (and more generally, mesenchymal cells). Our model assumptions to capture mesenchymal (not amoeboid) movement suggest that the lamellipodium plays a role in mediating cell-cell and cell-matrix adhesions and, ultimately, in producing the distinct behavior of frontal traction adhesion. These differences in adhesion and motility strategies reflect the divergent evolutionary adaptations of Dd and MDCK cells, with amoeboid migration favoring rapid, flexible movement and mesenchymal migration relying on stable, adhesion-dependent protrusions. Despite the fundamental differences between MDCK and Dd cell types, pairs of MDCK cells migrate comparable distances to Dd cell pairs, albeit on longer timescales; MDCK pairs require several hours to traverse distances that Dd cells cover within minutes. It is worthwhile to mention that in our model the same parameters are used, specifically the same viscous drag coefficient, which results in the same timescale of the dynamics. There are a number of other possibilities that could account for the differences in the observed traction patterns of motile Dd and MDCK-like cells. Namely, that the polarization machinery is differently modulated during collective migration across cell types. One model predicts a spectrum of collective cell motility modes for MDCK cells on fibronectin tracks, based on the interplay between polarization activity and intracellular adhesion forces [61]. In agreement with the model proposed by Ron et al., we also find that the strength of cell-cell adhesion forces can alter the dominant migration mode in MDCK-like pairs (but not Dd pairs). Another possibility is due to differences in the properties (such as contractility) between cells in a pair. While cell-to-cell differences are not accounted for in this model due to the lack of experimental data to support this assumption, previous work from our group has looked at the molecular requirements to capture the supracellular morphology of doublets based on experimental data from Ciona heart progenitor cells *Ciona* heart progenitor cells [18]. Lastly, other, more complex models have been proposed to capture other aspects of *in vitro* findings. For example, Rossetti et al. use an active gel theory model to show that migration efficiency decreases with cell chain length, an experimentally observed feature that our model does not capture [26].

There is much complexity of collective cell chemotaxis that our study did not address. Complex, interdependent molecular pathways carefully regulate polarization activity in response to chemical cues, along with cell-cell and cell-matrix adhesions. A careful analysis should incorporate a cytoskeletal network with these molecular actions of protrusive activity and adhesion complexes anchored not just to the underlying matrix, but also to the cytoskeleton. Indeed, in our formulation, protrusive activity is simply constant in time and in a fixed direction. Just as in our single cell model, one emergent characteristic of this model is the localization of tangential traction adhesions, but during any particular phase of the motility cycle we have three or four distinct adhesion sites rather than the two or three as reported experimentally. We attribute this difference to the simplified model of the actin cytoskeleton; here, the cytoskeleton only transmits the leading-edge forces to the surface on the entire region of contact. A more explicit description of the cytoskeleton should capture the non-uniform transmission of forces. Other sources of complexity not addressed by this work are natural cell-to-cell variability, and stochastic effects in collective chemotaxis. Another limitation stems from the restriction to two dimensions – lateral stresses are not available in this formulation and, indeed, the distribution between lateral and axial stresses is believed to play a crucial role in collective migration efficiency [62]. Despite these limitations, our model is not only able to capture various motility modes and their biased distributions observed in two different experiments but also provides a fundamental framework to understand collective migratory behavior across cell types and scales. Importantly, the work highlights the role of membrane protrusions in regulating cell-cell and cell-matrix adhesions, and ultimately, determining the dominant motility mode. Future research will shed light on the impact of these complexities on the emergent motility modes in collective cell migration.

## ACKNOWLEDGMENTS

We thank Sam Walcott for helpful discussions. We are grateful to Libera Lo Presti for revising the manuscript. We acknowledge that part of the experimental data of *Dictyostelium discoideum* migrating cell pairs originates from [8] and were adapted in accordance with the license terms of the Molecular Biology of the Cell journal. This work was supported in part by NSF DMS2209494 (C.C.), and the Deutsche Forschungsgemeinschaft (DFG) under Germany’s Excellence Strategy – EXC 2124 – 390838134 (E.E.B). This work was completed in part using the Discovery cluster, supported by Northeastern University’s Research Computing team.

